# Neutralization of Omicron SARS-CoV-2 by 2 or 3 doses of BNT162b2 vaccine

**DOI:** 10.1101/2022.01.21.476344

**Authors:** Hongjie Xia, Jing Zou, Chaitanya Kurhade, Hui Cai, Qi Yang, Mark Cutler, David Cooper, Alexander Muik, Kathrin U. Jansen, Xuping Xie, Kena A. Swanson, Pei-Yong Shi

## Abstract

We report the antibody neutralization against Omicron SARS-CoV-2 after 2 and 3 doses of BNT162b2 mRNA vaccine. Vaccinated individuals were serially tested for their neutralization against wild-type SARS-CoV-2 (strain USA-WA1/2020) and an engineered USA-WA1/2020 bearing the Omicron spike glycoprotein. Plaque reduction neutralization results showed that at 2 or 4 weeks post-dose-2, the neutralization geometric mean titers (GMTs) were 511 and 20 against the wild-type and Omicron-spike viruses, respectively, suggesting that two doses of BNT162b2 were not sufficient to elicit robust neutralization against Omicron; at 1 month post-dose-3, the neutralization GMTs increased to 1342 and 336, respectively, indicating that three doses of vaccine increased the magnitude and breadth of neutralization against Omicron; at 4 months post-dose-3, the neutralization GMTs decreased to 820 and 171, respectively, suggesting similar neutralization decay kinetics for both variants. The data support a three-dose vaccine strategy and provide the first glimpse of the neutralization durability against Omicron.

## Main text

Severe acute respiratory syndrome coronavirus 2 (SARS-CoV-2) continues to evolve and generates new variants of concern (VOCs), including Alpha, Beta, Gamma, Delta, and Omicron. Variant mutations, particularly those in the spike glycoprotein, can alter viral transmission, disease severity, therapeutic antibody efficacy, and potentially vaccine effectiveness. The newly emerged Omicron variant (B.1.1.529 lineage) was first detected in South Africa, reported to WHO on November 24^th^, 2021, and designated as a VOC two days later. It is rapidly spreading around the world and causing increased breakthrough infections in previously infected and vaccinated individuals.^1^ Laboratory results are urgently needed to examine vaccine-elicited neutralization of the Omicron variant and the duration in which this activity persists.

BNT162b2, an mRNA vaccine that encodes the prefusion stabilized full spike glycoprotein of SARS-CoV-2 isolate Wuhan-Hu-1, has received regulatory approval and/or authorization around the world including approval for vaccination of people 16 years of age and older and authorization under emergency use provision for immunization of individuals 5-15 years old by the US Food and Drug Administration. We formerly reported that recombinant SARS-CoV-2 bearing spike genes from previous VOCs, variants of interest (VOIs), and variants under monitoring (VUMs) remained susceptible to BNT162b2 vaccine-elicited neutralization. While the Beta variant exhibited the most neutralization reduction among all tested variants, high efficacy against Beta was observed.^2^

To determine the susceptibility of Omicron variant to BNT162b2-elicited neutralization, we engineered the complete Omicron *spike* into the USA-WA1/2020 genetic background (**Fig. S1**). The resulting recombinant Omicron-spike SARS-CoV-2 contained spike mutations A67V, H69-V70 deletion (Δ69-70), T95I, G142D, V143-Y145 deletion (Δ143-145), N211 deletion (Δ211), L212I, L214 insertion EPE (ins214EPE), G339D, S371L, S373P, S375F, K417N, N440K, G446S, S477N, T478K, E484A, Q493R, G496S, Q498R, N501Y, Y505H, T547K, D614G, H655Y, N679K, P681H, N764K, D796Y, N856K, Q954H, N969K, and L981F (*GISAID EPI_ISL_6640916*). The recombinant Omicron-spike SARS-CoV-2 and wild-type USA-WA1/2020 produced infectious titers of greater than 10^7^ plaque-forming units per milliliter (PFU/ml). The recombinant viruses were sequenced to ensure no undesired mutations. Although Omicron-spike virus formed smaller plaques than the wild-type USA-WA1/2020 (**Fig. S2**), both viruses showed equivalent viral RNA genome/PFU ratios when analyzed on Vero E6 cells (**Fig. S3**), suggesting comparable specific infectivities of the viral stocks.

We determined 50% plaque reduction neutralization titers (PRNT_50_) against recombinant USA-WA1/2020 and Omicron-spike SARS-CoV-2 using four serum panels from BNT162b2-vaccinated participants of the phase 1 portion of Study C4591001. The first panel contained 20 sera, collected 2 or 4 weeks after the second dose of 30 μg of BNT162b2, which was administered 3 weeks after the first immunization^2^ (**Fig. S4**). These post-dose-2 (PD2) sera neutralized USA-WA1/2020 and Omicron-spike SARS-CoV-2 with geometric mean titers (GMTs) of 511 and 20, respectively. All PD2 sera had neutralization titers of 160 or higher against the wildtype USA-WA1/2020, whereas only 11 of the 20 sera had neutralization titers of 20 (limit of detection) or higher against the Omicron-spike virus (**Fig. 1A**). The second serum panel contained 22 specimens, collected on the day of the third 30 μg BNT162b2 dose which was administered 7.9 to 8.8 months after dose 2^3^ (**Fig. S4**). The GMTs against USA-WA1/2020 decreased to 65, while the GMTs against Omicron remained low at 13 (**Fig. 1B**). The third and fourth serum panels were collected 1 and 4 months after the third dose, respectively (**Fig. S4**). The 1-month-post-dose-3 (1MPD3) sera neutralized USA-WA1/2020 and Omicron-spike SARS-CoV-2 with GMTs of 1342 and 336, respectively (**Fig. 1C**). The 4-month-post-dose-3 (4MPD3) sera neutralized USA-WA1/2020 and Omicron-spike SARS-CoV-2 with GMTs of 820 and 171, respectively; all the 4MPD3 sera neutralized Omicron-spike virus at titers of 28 or higher (**Fig. 1D**). **Figure S5** summarizes the PRNT_50_ curves for individual subjects. Collectively, the data support three conclusions. First, at 2 or 4 weeks PD2, the neutralization GMT against Omicron-spike SARS-CoV-2 was 25.6 times lower than that against USA-WA1/2020 and in a similar range of reduction noted in some preliminary and peer reviewed reports.^4^ Differences in Omicron neutralization reduction compared to wild-type USA-WA1/2020 across studies are likely attributed to different serum samples, time of sampling, assay formats and assay protocols, but nonetheless support a substantial decrease in neutralization activity against Omicron after two doses of BNT162b2. Second, by 1-month PD3, neutralization GMTs against wild-type USA-WA1/2020 and Omicron-spike virus increased by 2.6 and 16.8 times, respectively, when compared to the corresponding GMTs by 2 to 4 weeks PD2. Thus, after dose 3, the increase of neutralization GMTs against the Omicron virus was 6.5 times higher than that against USA-WA1/2020. This result agrees with our recent findings that a third dose of BNT162b2 increased the magnitude and breadth of neutralization against Delta and Beta variants.^3^ Third, from 1 to 4 months PD3, neutralization GMTs against USA-WA1/2020 and Omicron decreased by 1.6 and 2 times, respectively, suggesting similar antibody decay kinetics for both variants.

**Figure 1.**
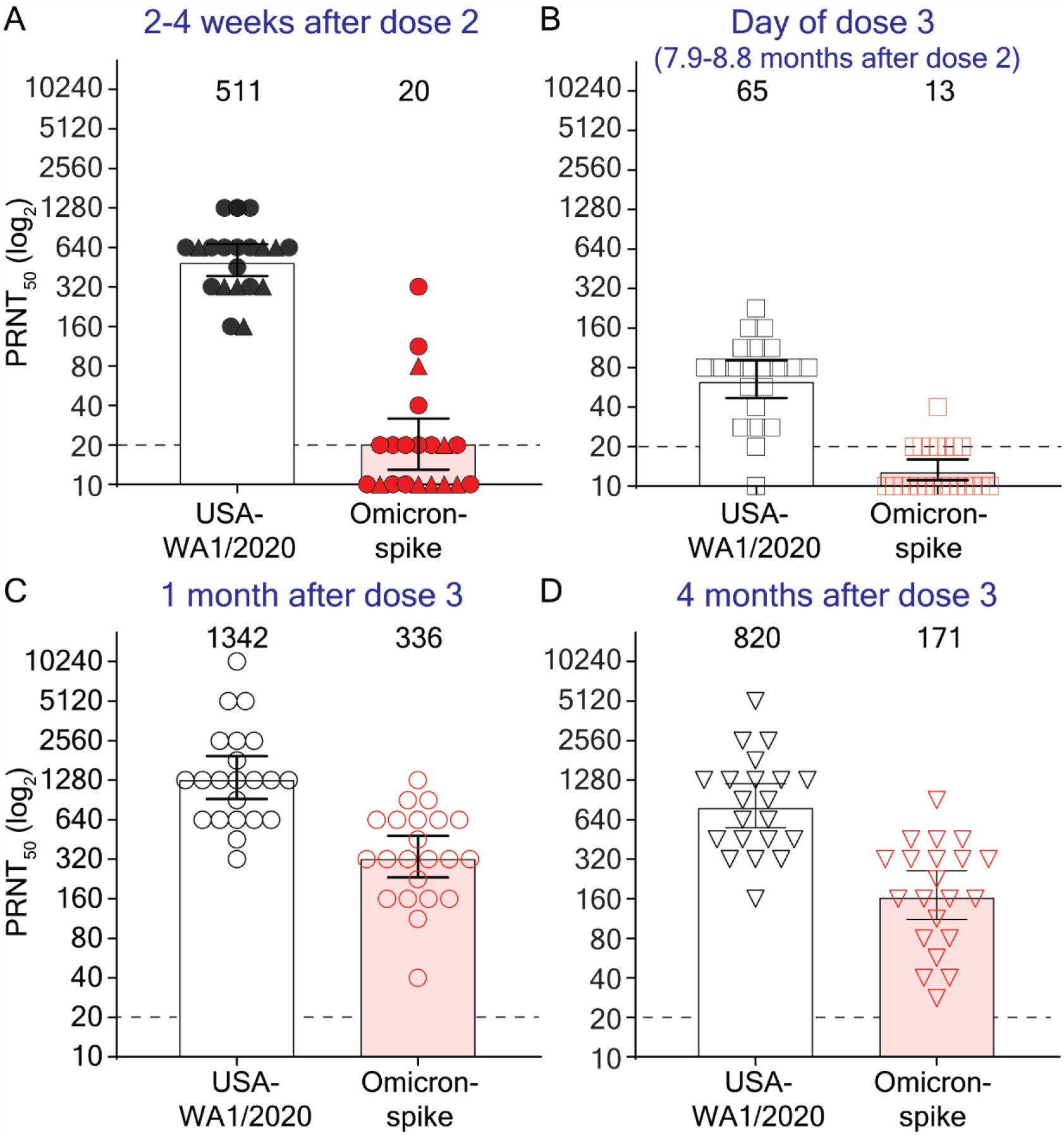
Serum neutralization of USA-WA1/2020 and Omicron-spike SARS-CoV-2 after two and three doses of BNT162b2. Shown are the 50% plaque reduction neutralization titers (PRNT_50_) for four panels of BNT162b2-vaccinated human sera. The first panel contained 20 specimens obtained at 2 weeks (circles) or 4 weeks (triangles) after the second dose of BNT162b2 (**A**). The second panel contained 22 specimens collected on the day of the third dose of BNT162b2 (**B**), which was administered at 7.9 to 8.8 months after the second dose. The third panel contained 22 specimens collected at one month after the third dose (**C**). The fourth panel contained 21 specimens collected at 4 months after the third dose (**D**). The information for the first and third serum panels were reported previously.^2,3^ The Omicron-spike SARS-CoV-2 was produced by engineering the complete Omicron spike gene into USA-WA1/2020. Each data point represents the geometric mean PRNT50 obtained with a serum specimen against the indicated virus, as detailed in **Table S1**. The neutralization titer was determined in duplicate assays, and the geometric mean was presented. The bar heights and the numbers above indicate geometric mean titers. The whiskers indicate 95% confidence intervals. The dotted line indicates the limit of detection of PRNT_50_. PRNT_50_ of < 20 was treated as 10 for plot purposes and statistical analysis. Statistical analysis was performed with the use of the Wilcoxon matched-pairs signed-rank test. The statistical significance of the difference between geometric mean titers in the USA-WA1/2020 and Omicron-spike SARS-CoV-2 is p < 0.0001.

Our data demonstrate that three doses of BNT162b2 elicited substantial neutralization activity against Omicron-spike SARS-CoV-2, while two doses showed significantly reduced neutralization titers. The neutralization GMT against Omicron-spike virus after dose 3 (336) was close to the neutralization GMT against wild-type USA-WA1/2020 after dose 2 (511), which was previously associated with high efficacy in the pivotal efficacy study (C4591001). These data suggest that a third-dose vaccine strategy could minimize the health impact of Omicron. This is supported by initial reports demonstrating that while vaccine effectiveness against symptomatic infection due to Omicron may wane following a third dose of BNT162b2, the effectiveness against hospitalization remains high.^5^

The current study has two limitations. First, the durability of the neutralization against Omicron variant beyond 4 months PD3 remains to be determined. Second, other immune effectors, such as T cells and non-neutralizing antibodies that mediate antibody-dependent cytotoxicity, have not been examined. CD8+ T cell responses likely play an important role in protection against severe COVID-19; despite the significant number of mutations in the Omicron spike, the majority of T cell epitopes of spike glycoprotein are preserved. Taken together, the current in vitro study suggests that 2 doses of BNT162b2 may remain effective against severe disease, but that a third dose of BNT162b2 is likely required to maintain effectiveness against COVID-19 caused by Omicron. Additional real world effectiveness data and laboratory investigations will further inform the duration of protection, potential need for an additional dose at a later time, and whether an Omicron modified vaccine is required.

## Materials and Methods

### Construction and characterization of recombinant Omicron-spike SARS-CoV-2

Recombinant Omicron-spike SARS-CoV-2 was constructed by engineering the complete spike gene from Omicron variant (GISAID EPI_ISL_6640916) into an infectious cDNA clone of clinical isolate USA-WA1/2020^6^ **(Figure S1**). All spike mutations were introduced into the infectious cDNA clone of USA-WA1/2020 using PCR-based mutagenesis as previously described.^7^ The full-length cDNA of viral genome containing the complete Omicron spike mutations was assembled via *in vitro* ligation. The resulting full-length cDNA was used as a template for *in vitro* transcription of genome-length viral RNA. The *in vitro* transcribed genome-length viral RNA was electroporated into Vero E6 cells. On day 3 post electroporation, the original viral stock (P0) was harvested from the electroporated cells. The P0 virus was propagated for another round on Vero E6 cells to produce the P1 stock for neutralization testing. The infectious titer of the P1 virus was quantified by plaque assay on Vero E6 cells (**Figure S2**). The complete spike gene of the P1 virus was sequenced to ensure no undesired mutations. The protocols for the mutagenesis of SARS-CoV-2 and virus production were reported previously.^8^ For determining the specific infectivity of each virus, the P1 virus stock was quantified for its plaque-forming unit (PFU) and genomic RNA content by plaque assay on Vero E6 cells and RT-qPCR, respectively. The methods for plaque assay and RT-qPCR were reported previously.^9^ The specific infectivity of each virus was indicated by the genomic RNA-to-PFU ratios (genome/PFU).

### Serum samples and plaque-reduction neutralization testing

Four BNT162b2-vaccinated serum panels were used in the study. **Figure S4** summarizes the collection time of serum panels. The first panel (n=20) was collected at 2 weeks (circles) or 4 weeks (triangles) after the second dose of BNT162b2. The second panel (n=22) was collected on the day of the third dose of BNT162b2, which was administered at 7.9 to 8.8 months after the second dose. The third (n=22) and fourth (n=21) panels were collected at 1 and 4 months after the third dose, respectively. The information for the first and third serum panels were reported previously.^2,3^ The 50% plaque-reduction neutralization titer (PRNT_50_) was measured for each serum as previously reported.^10,11^ Individual sera were 2-fold serially diluted in culture medium with a starting dilution of 1:20. The diluted sera were incubated with 100 PFU of Omicron-spike SARS-CoV-2 or USA-WA1/2020. After 1 h incubation at 37 °C, the serum-virus mixtures were inoculated onto 6-well plates with a monolayer of Vero E6 cells pre-seeded on the previous day. The PRNT_50_ value was defined as the minimal serum dilution that suppressed >50% of viral plaques. The neutralization titer was determined in duplicate assays, and the geometric mean was taken. **Table S1** summarizes the PRNT_50_ results for all serum samples.

## Acknowledgments

We thank the Pfizer-BioNTech clinical trial C4591001 and NCT04368728 participants, from whom the post-immunization human sera were obtained. We thank the many colleagues at Pfizer and BioNTech who developed and produced the BNT162b2 vaccine. We thank John Yun-Chung Chen and Cody Bills from the University of Texas Medical Branch at Galveston for help with plaque assays.

---

**Figure S1.**
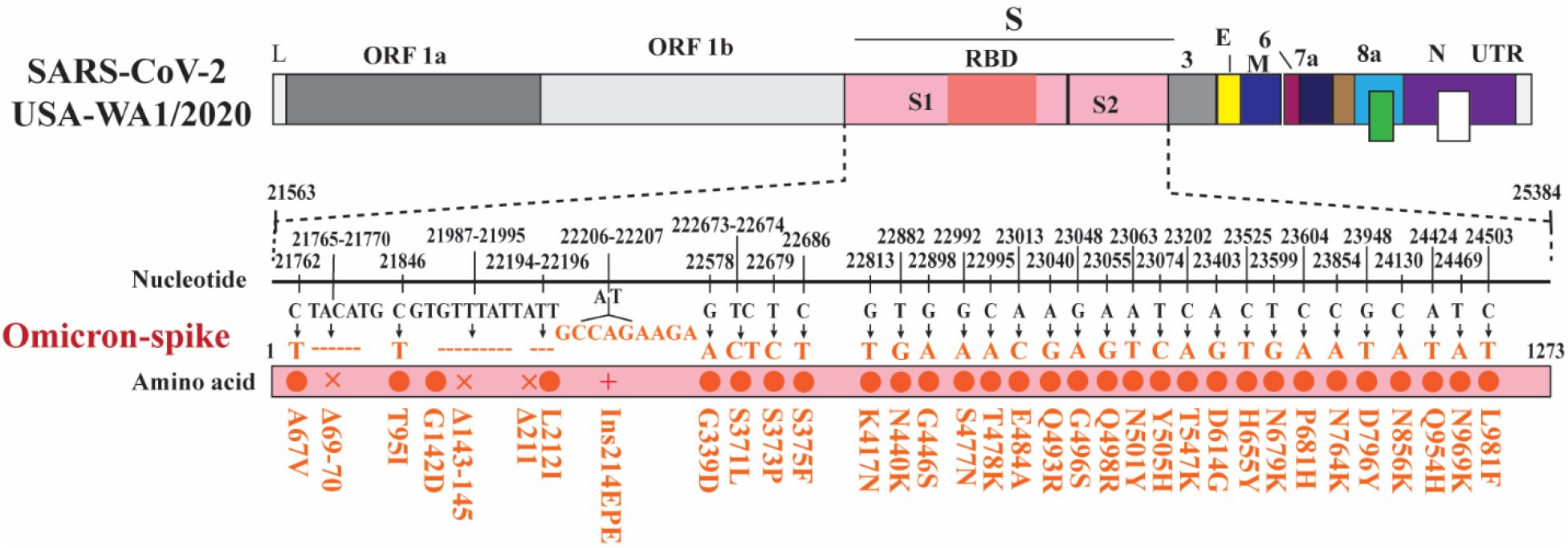
Diagram of engineered Omicron-spike SARS-CoV-2. USA-WA1/2020 was used as the wild-type virus to engineer Omicron-spike SARS-CoV-2. Mutations (red circle), deletions (x), and insertions (+) are indicated. Nucleotide and amino acid positions are depicted. L: leader sequence; ORF: open reading frame; RBD: receptor binding domain; S: spike glycoprotein; S1: N-terminal furin cleavage fragment of S; S2: C-terminal furin cleavage fragment of S; E: envelope protein; M: membrane protein; N: nucleoprotein; UTR: untranslated region.

**Figure S2.**
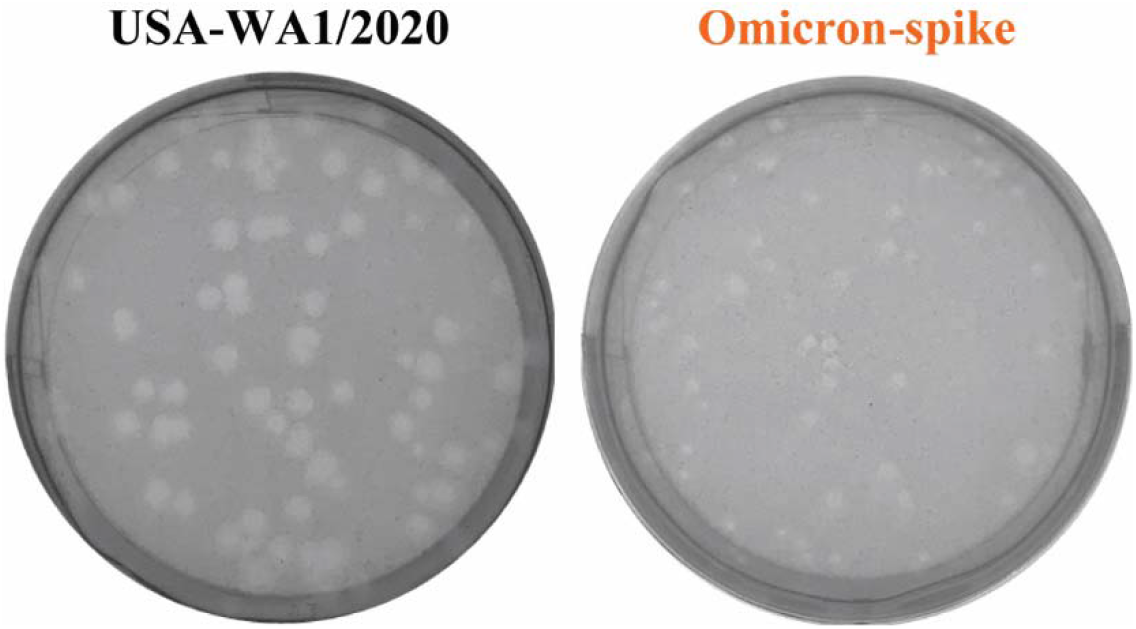
Plaque morphologies of USA-WA1/2020 and Omicron-spike SARS-CoV-2. The plaque assay was performed on Vero E6 cells in 6-well plates.

**Figure S3.**
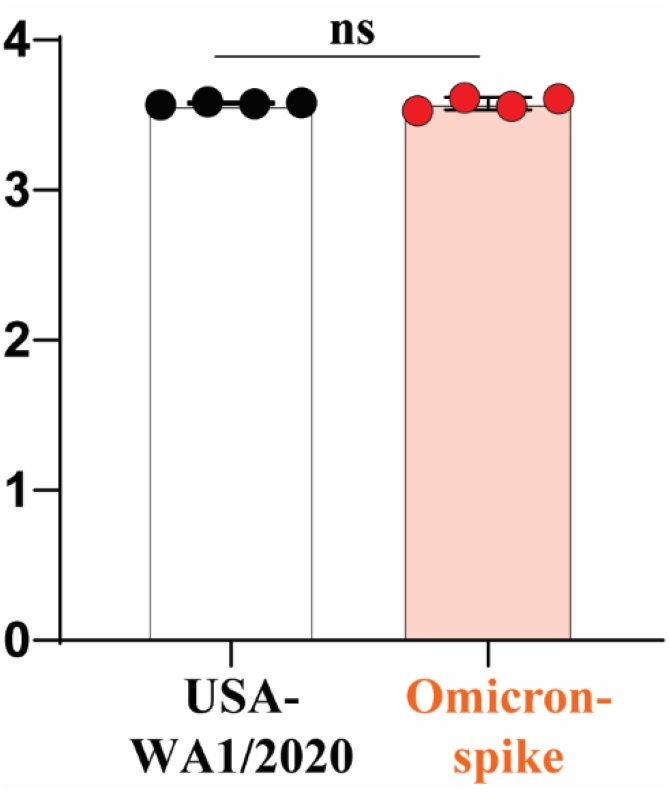
Viral genomic RNA versus plaque-forming unit ratios (genomes/PFU) of USA-WA1/2020 and Omicron-spike SARS-CoV-2. Individual recombinant virus P1 stocks were measured for their viral RNA and plaque-forming units by RT-qPCR and plaque assay on Vero E6 cells, respectively. The specific infectivity of each recombinant virus was determined by the viral genome/PFU ratio. Dots represent individual biological replicates of 4 different aliquots of viruses. The values in the graph represent the means with 95% confidence intervals. A nonparametric Mann-Whitney test was used to determine significant difference between USA-WA1/2020 and Omicron-spike SARS-CoV-2. n.s., no statistical difference.

**Figure S4.**
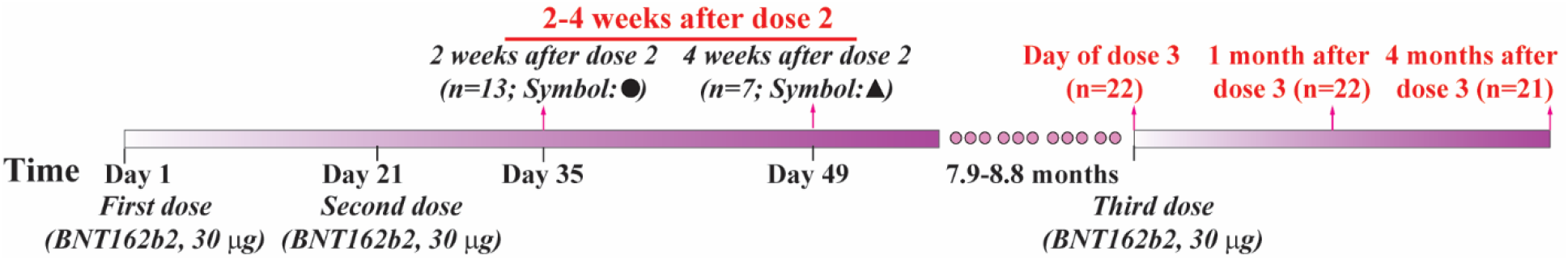
BNT162b2-vaccinated serum collection. Four BNT162b2-vaccinated serum panels were used in the study. The first panel had 20 samples obtained at 2 weeks (circles) or 4 weeks (triangles) after the second dose of BNT162b2. The second panel (n=22) was collected on the day of the third dose of BNT162b2. The third (n=22) and fourth (n=21) panels were collected at 1 and 4 months after the third dose, respectively. The information for the first and third serum panels were reported previously.^2,3^

**Figure S5.**
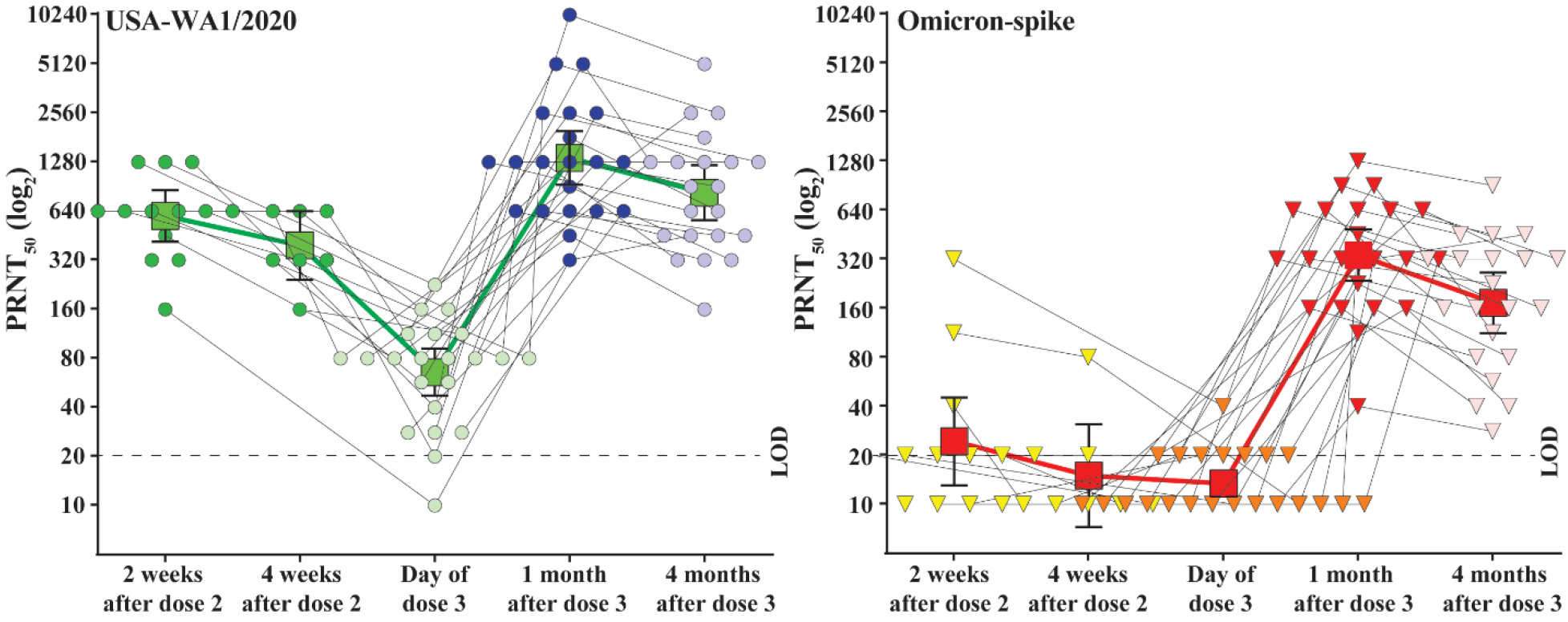
Neutralization of BNT162b2-vaccinated sera against USA-WA1/2020 and Omicron-spike SARS-CoV-2. The PRNT_50_ data from individual subjects are connected with lines. The GMTs for each time point are presented as squares. Error bars indicate the 95% CI of the GMTs. The dotted lines indicate the limit of detection (LOD) of PRNT_50_. PRNT_50_ of < 20 was plotted as 10. The original PRNT_50_ values are presented in **Table S1**.

**Table S1.**
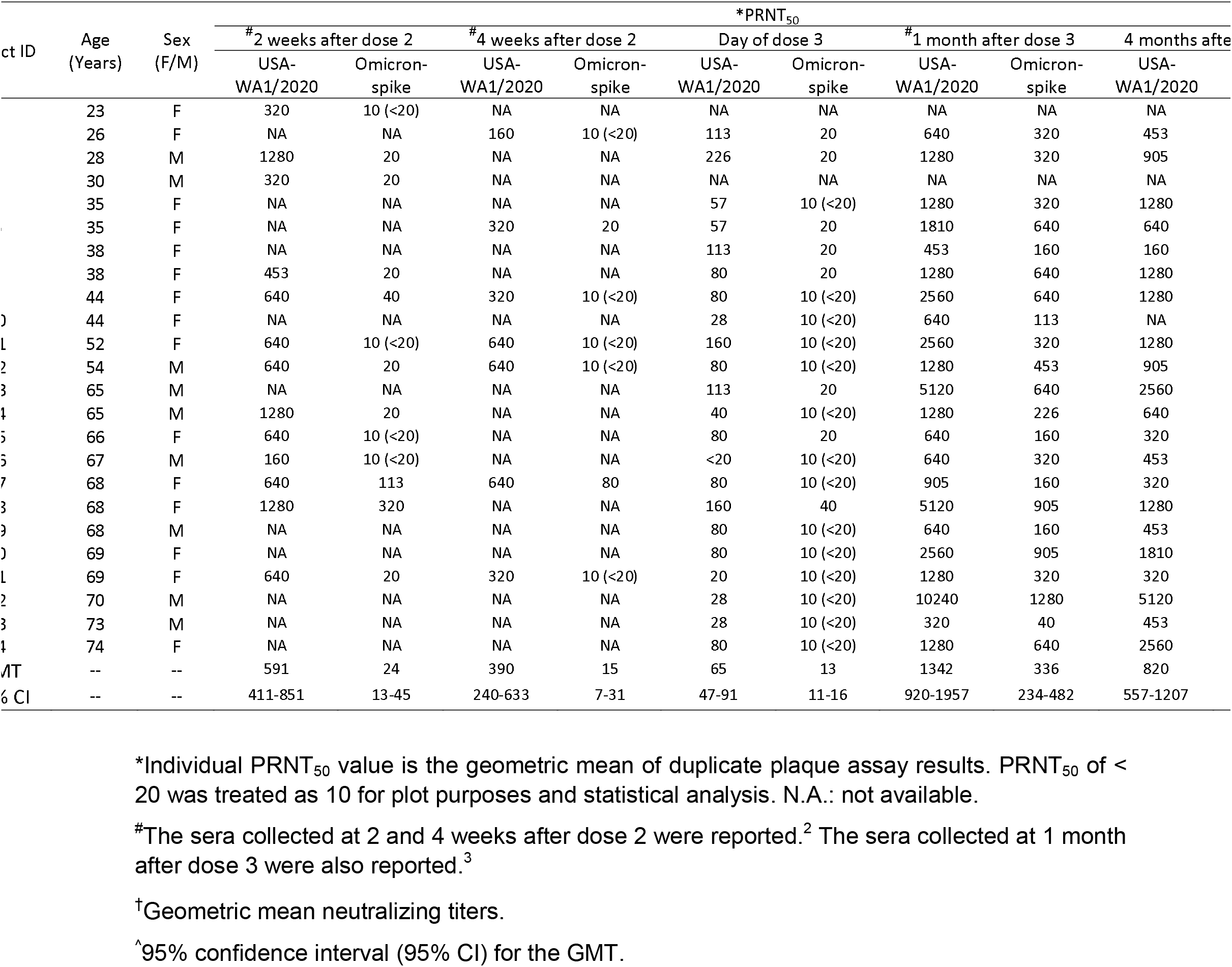
PRNT_50_ values of BNT162b2-vaccinated sera against USA-WA1/2020 and Omicron-spike SARS-CoV-2.

